# M2M-InvNet: Human Motor Cortex Mapping from Multi-Muscle Response Using TMS and Generative 3D Convolutional Network

**DOI:** 10.1101/2022.07.22.501062

**Authors:** Md Navid Akbar, Mathew Yarossi, Sumientra Rampersad, Kyle Lockwood, Aria Masoomi, Eugene Tunik, Dana Brooks, Deniz Erdoğmuş

**Author notes:** Corresponding author: Md Navid Akbar. This project was funded in part by grants NIH R01NS085122 and R01HD058301 (ET), NSF CMMI-193537 (ET,DE,MY), NSF CBET-1804550 (ET,DE,DB), NSF CBET-2117626 (ET,DE,DB,MY,SR).

## Abstract

Transcranial magnetic stimulation (TMS) is often applied to the motor cortex to stimulate a collection of motor evoked potentials (MEPs) in groups of peripheral muscles. The causal interface between TMS and MEP is the selective activation of neurons in the motor cortex; moving around the TMS ‘spot’ over the motor cortex causes different MEP responses. A question of interest is whether a collection of MEP responses can be used to identify the stimulated locations on the cortex, which could potentially be used to then place the TMS coil to produce chosen sets of MEPs. In this work we leverage our previous report on a 3D convolutional neural network (CNN) architecture that predicted MEPs from the induced electric field, to tackle an inverse imaging task in which we start with the MEPs and estimate the stimulated regions on the motor cortex. We present and evaluate five different inverse imaging CNN architectures, both conventional and generative, in terms of several measures of reconstruction accuracy. We found that one architecture, which we propose as M2M-InvNet, consistently achieved the best performance.

## I. Introduction

Transcranial magnetic stimulation (TMS) is a non-invasive technique that uses magnetic fields to stimulate neurons in the brain [1]. When TMS is applied to the motor cortex, it may result in muscle activation. This activation can be measured as motor evoked potentials (MEPs) using standard surface electromyography (EMG). By varying coil position over the motor cortex, TMS can be used non-invasively in humans as a causal probe to investigate the spatial topography of muscle activation patterns [2]. TMS mapping of cortical muscle topography has shown clinical utility [3], for example, to quantify cortical muscle topography associated with abnormal muscle activation patterns due to stroke and track changes during recovery [4], [5], and to perform the presurgical evaluation of motor, speech, or language functions for patients requiring resections in eloquent areas [6], [7]. Advances in modeling of the TMS-induced E-field [8], [9] have allowed greater resolution in the estimation of the cortical representations underlying evoked muscle activation. These approaches link information about the induced electric fields in the cortex to the stimulation intensity and orientation dependent responses in single muscles [9], [10]. Recently, work from our group has proposed that TMS may be used to study patterns of multi-muscle activation that have been theorized to form the basis of modular control of coordinated movement [11], [12].

Previously, we developed a forward model using a convolutional neural network (CNN) autoencoder (AE) and a separate deep CNN mapper that connects the simulated E-field and recorded MEPs to estimate multi-muscle activation patterns induced by new TMS stimulations [11], [12]. To our knowledge, this was the first report of a robust computational *forward modeling* framework going from TMS-induced E-Fields to multi-muscle MEPs. In the present study, we expand on our previous forward modeling technique by developing an *inverse modeling* approach to estimate (putatively causal) cortical E-fields from muscle activation patterns recorded from a collection of relevant muscles. In other words, our system can predict which region of the motor cortex was stimulated by the TMS coil based on a multi-muscle MEP pattern. The proposed model is intended to be subject-specific, targeting applications such as preoperative mapping and tracking recovery, where patient-specific data is required. Furthermore, the model offers a potential tool to investigate cortical representations of motor modularity non-invasively and may, pending future validation, be applied to clinical populations as a diagnostic tool to explain cortical contributions to pathological movement patterns.

We start with subject-specific volume conduction models based on magnetic resonance images (MRIs), followed by finite element (FE) modeling of the E-fields based on the position and orientation of the TMS coil. We report on five deep network architectures that were developed based on selected combinations of CNNs and variational inference (VI). We chose these tools because CNNs have previously been used for TMS modeling to generate head models [13], and to estimate induced E-fields directly from MRI scans [14]. In addition, CNN AEs using VI, known as variational autoencoders (VAEs), have been widely used in computer vision for natural-looking image reconstruction, since deep generative models such as a VAE can constrain the reconstructed image to remain on a learned underlying manifold, such that the reconstructions are more physically or biologically meaningful [15]. VAEs have also matched the performance of standard compressed sensing techniques in inverse imaging with less training data [16]. Three of the five models we developed utilized a two-stage training strategy [16]: first learning a latent space from the E-fields, and second refining that space by learning from the MEP mapping. The remaining two models jointly learned the latent space from the MEPs and the E-fields in a single-stage training strategy [17].

To carry out our study, we collected MRI scans, TMS coil position and orientation, and 15-muscle MEP data from three healthy subjects during expert user-guided cortical motor topography mapping. We stimulated at *∼*1,000 scalp locations per subject (699, 1200, and 1199 for subjects 1, 2 and 3, respectively). We used a stratified train-validate-test cross-validation approach to evaluate the ability of each of these five networks to accurately estimate the stimulated cortical region, as determined by the FE modeling, that produced a given MEP pattern.

Our results suggest that our networks can indeed perform this task with reasonable accuracy and robustness as long as there is sufficient MEP activity. The model that directly learns from cortical stimulation and MEPs jointly achieved the lowest squared error and the highest fidelity to reconstruction, across all subjects.

## II. Methods

All protocols were conducted in conformance with the Declaration of Helsinki and were approved by the Institutional Review Board of Northeastern University (IRB# 15-10-22, last approved September 23, 2021). Three healthy subjects (3 males, ages 25, 35, & 36) participated after providing institutionally approved written informed consent. All subjects were right-hand dominant according to the Edinburgh handedness inventory [18], free of neurological or orthopedic conditions that could interfere with the experiment, and met inclusion and exclusion criteria to receive TMS [19].

### A. Data Acquisition

The procedure used for TMS mapping has been previously described in detail [11]. Briefly, subjects were seated comfortably with the right upper limb supported in an arm trough, and the left upper limb resting comfortably on an armrest. Surface EMG (Delsys Inc., Natick, MA) was recorded at 2000 Hz (common mode rejection ratio *>*80 dB, 99.99% Ag, built-in 20–450 Hz bandpass filter) from 15 hand and arm muscles: 1st dorsal interosseus (FDI), 3rd dorsal interosseous (3DI), 3rd lumbrical (3Lum), extensor indicus (EI), abductor pollicis brevis (AbPB), adductor pollicis brevis (AdPB), abductor digiti minimi (ADM), flexor digiti minimi (FDM), flexor carpi radialis (FCR), flexor carpi ulnaris (FCU), flexor digitorum superficialis (FDS), extensor digitorum (EDC), and extensor carpi radialis (ECR), extensor carpi ulnaris (ECU), brachioradialis (BRD). Delsys Trigno Mini sensors specialized for the measurement of small muscles were used for the collection of all intrinsic hand muscles (FDI, EI, 3Lum, EI, AbPB, AdPB, and ADM). Standard Delsys Trigno sensors were used for all remaining muscles tested. Care was taken in electrode placement to limit the potential for cross-talk between sensors.

To ensure spatial TMS precision, frameless neuronavigation (Brainsight, Rogue Research) was used to co-register each subject’s head position with a 3D cortical surface rendering of their high-resolution anatomical MRI scan (T1-weighted, TI = 1100 ms, TE = 2.63 ms, TR = 2000 ms, 256*×*192*×*160 acquisition matrix, 1 mm^3^ voxels). TMS was performed using a Magstim BiStim2 stimulation unit (The Magstim Company Ltd) which delivers a monophasic pulse (*∼*100 *µ*s rise time, 1 ms pulse duration). The TMS coil (Magstim D702 70 mm figure-of-eight coil, monophasic pulse) was held tangential to the scalp with the handle posterior 45^*°*^ off the sagittal plane inducing a posterior-anterior current in the brain [20]. Motor evoked potentials were measured as the peak-to-peak EMG amplitude 10-50 ms after the TMS pulse [5], [11], [21]. The FDI muscle hotspot was found via a coarse map of the hand knob area to identify the location that produced the largest and most consistent MEP amplitudes [5], [22], [23]. Resting motor threshold (RMT) was selected as the minimum intensity required to elicit MEPs *>*50 *µ*V on 3 out of 6 consecutive stimulations [5]. In a single experimental session, TMS maps were collected at stimulus intensities of 110%, 120%, 130%, and 140% of RMT. The distribution of both the number of stimulations chosen after preprocessing and those originally applied, corresponding to each map for each subject, is reported in Table I. The details of these preprocessing techniques are outlined in Section II-C. For each map, TMS (100-300 stimulations, 4-5 jittered ISI) was delivered along the vertices of a 6×6 cm regular grid (1 cm spacing, 36 cm^2^ area, 7×7=49 vertices) centered on the hotspot. For each intensity, one stimulus was delivered to each of the 49 equidistant points on the predefined grid. The remaining stimuli (51-251 per intensity for subject 1 and 250-251 per intensity for subjects 2 and 3) were delivered within the 6×6 cm area defined by the grid at loci selected by the expert TMS operator using real-time feedback from the MEPs to maximize information about the responsive areas. We have previously shown that this technique produces similar information to traditional gridded mapping approaches [24]. Care was taken to ensure that the mapping included the full extent of the excitable area at the given stimulation intensities for all recorded muscles. For each pulse, MEP amplitudes were recorded of the 15 muscles selected for analysis.

**TABLE I.** Distribution of both the number of stimulations chosen after preprocessing and those originally applied, for each stimulation intensity in each subject.

| Subject | % of RMT |  |  |  |
| --- | --- | --- | --- | --- |
|  | 110 | 120 | 130 | 140 |
| 1 | 196/300 | 63/149 | 119/150 | 87/100 |
| 2 | 298/300 | 299/300 | 293/300 | 285/300 |
| 3 | 299/300 | 281/299 | 282/300 | 292/300 |

\* The numbers are represented in a p/q format, where q indicates the total number of stimulations applied and p indicates the actual number chosen after preprocessing.

### B. Finite Element Modeling of TMS E-fields

The model for subject 1 was constructed as described in [11] and the simulation data from that study was used for the analyses in this paper. The models for subjects 2 and 3 were constructed specifically for this work. For these models, individual T1 MRI scans (TR = 1.9 ms, TE = 2.0 ms, 256*×*256*×*176 mm, 1 mm^3^ voxels) were processed by the SimNIBS headreco algorithm [25] to produce tissue segmentations of skin, bone, skull cavities, eyes, cerebrospinal fluid, gray and white matter. Segmentation masks were manually corrected in Corview (MARREK Inc., Salt Lake City, UT), converted to surface meshes by headreco, and combined into a tetrahedral volume mesh with TetGen [26], resulting in meshes with 3.6 and 3.8 million elements. Isotropic conductivity values were assigned to each element based on tissue type [27]. The TMS coil was modeled in SCIRun [28] using the BrainStimulator toolbox [29] by approximating the coil field as the magnetic vector potential from small magnetic dipoles distributed across the coil [30]. Simulation of electric fields induced by this coil in the head model was conducted using a quasi-static FE framework implemented in BrainStimulator [29]. The resulting E-field for each coil position was then spatially resampled to a hexahedral mesh (1 mm^3^ elements). Using Freesurfer [31], the cortex on the MRI scan was segmented into Brodman areas. The mask for area BA4 was used to select the corresponding voxels in the hexahedral mesh. A 64×64×64 section of the grid surrounding the BA4 area was used as input for the next stage of the modeling procedure.

### C. Preprocessing

Several steps were taken to clean and prepare the data. First, some of the stimulations did not produce any muscle activation. These stimulations were identified automatically (by checking if all the normalized MEP values were zero) and subsequently removed from further analysis to avoid null space problems for the network. Some outlier stimulations that resulted in unusually low E-field values in the BA4 area (maximum simulated E-field intensity *<*10 mV/m), which we believe were due to experimental errors, were also removed. These stimulations were only applicable for subject 1, and constituted of 49 out of the 149 total for 120% of RMT. Eventually, the actual number of stimulations chosen after applying these preprocessing steps are also reported in Table I.

For network training and testing, min-max scaling was used to preprocess the data. For the E-fields, the intensity of the voxel corresponding to the maximum strength of the E-fields in the entire set of stimulations for a particular subject was scaled to 1, voxels outside the BA4 motor cortex area were scaled to zero, and all other voxels were linearly scaled in that range. For the MEPs, each individual muscle activation was scaled to the unit interval [0, 1], with 1 representing the maximum activity of that muscle in the entire set of stimulations, for a given subject.

### D. Latent Variable Model

The causal forward model (E-field to MEP mapping) in [12] was expressed as

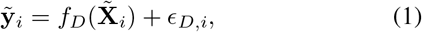

where **y**_*i*_ is the *m ×* 1 observed muscle activity vector (*m*=number of measured muscles) for the *i*-th stimulation, 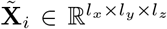 is the 3D E-field distribution on the motor cortex, *f*_*D*_() represents the CNN forward model (M2M-Net) [12] for direct mapping of cortical E-fields to MEPs, *ϵ*_*D,i*_ *∼ 𝒩* (**0**, *σ*^2^**I**) is the *m ×* 1 residual mapping error assumed to follow a white additive Gaussian distribution, and *i ∈ {*1, 2, …, *I}* represents the index of the train or test stimulation. In this work, *m* = 15 and *l*_*x*_ = *l*_*y*_ = *l*_*z*_ = 64. Since the objective of this work was to reuse a similar architecture as M2M-Net to reconstruct the E-field from the MEPs along an inverse imaging path, we sought to obtain an 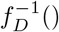 model such that

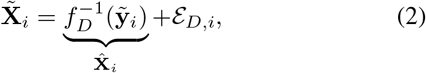

where 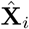 is the predicted E-field distribution, and 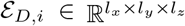 is the residual mapping error assumed to follow a zero mean, white additive Gaussian distribution.

Fig. 1 outlines the relations among the different variables, in the causal forward path and inverse imaging path. Here,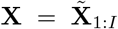 are the volumetric E-fields, 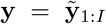 are the corresponding muscle activity vectors, and **z** is a *n ×* 1 latent variable vector, which is assumed to represent the individual subject’s cortico-motor mapping. The inverse imaging model 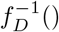, M2M-InvNet, consisted of a mapper block and a decoder block (both with and without an accompanying encoder). While in [12] only a standard AE was explored, four additional CNN architectures with and without VI were tested in this work, based on the idea that deep generative models (such as a VAE) might constrain **z** to remain on a learned manifold such that the reconstructions are more accurate [15].

**Fig. 1.**
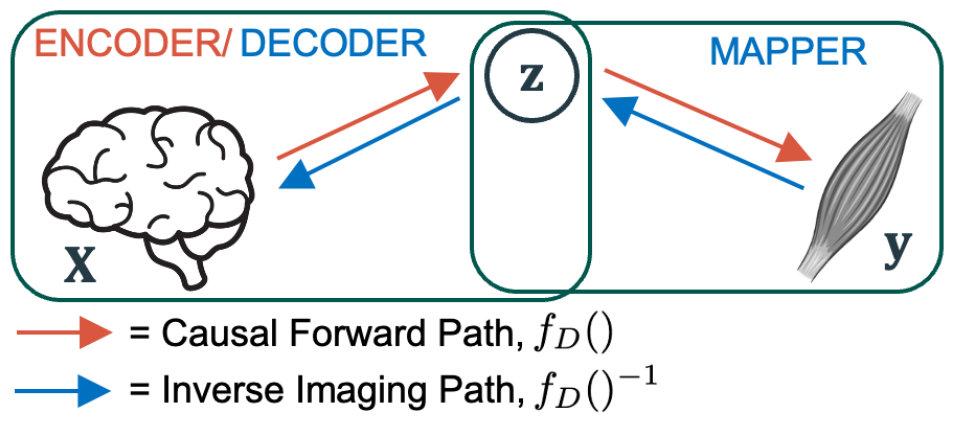
A high-level diagram of the system, outlining the relation of the observed **X** and **y** with the latent variable **z**, which represents the cortico-motor mapping. The red arrows indicate the path taken by the causal forward model, ***f***_***D***_(), which consists of the encoder (maps **z** from **X**). The blue arrows indicate the path taken by the inverse imaging model, ***f***_***D***_()^***−*1**^, which consists of the mapper (maps **z** from **y**) and the decoder (maps **X** from **z**). In testing mode only **y** is presented to the trained network and its task is to estimate **X**.

In this work, we designed five inverse models, namely: (a) AE-Decoder, (b) Direct Convolutional, (c) VAE-Decoder, (d) VAE-Sampler-Decoder, and (e) Direct Variational. Models (a), (c) and (d) utilize a two-stage training: they are first trained along a forward path (**X** *→* **z** *→* **y**), and then the learned **z** is utilized to guide the training in the reverse path (**y** *→* **z** *→* **X**), as seen in Fig. 1. In the forward path, model (a) uses an AE whereas models (c) and (d) use a VAE. Model (c) uses variational sampling only in the forward path, whereas model (d) loads the saved variational sampling from the forward path and re-trains it in the reverse path. Models (b) and (e) implement a single-stage training (**y** *→* **z** *→* **X**), with (b) using a purely convolutional architecture and (e) using a variational convolutional architecture.

Starting with the forward path in Fig. 1, the goal was to maximize the density function *P* (**X**) from a conditional distribution *P* (**X**|**z**) [32] as

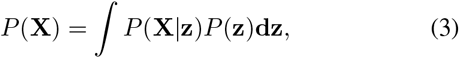

where **z** is to be sampled from the density function *P* (**z**). In a standard AE, *P* (**z**) is estimated by the encoder as *P* (**z**|**X**), whereas *P* (**X**|**z**) is approximated by the decoder. In a standard VAE, a surrogate distribution *Q*(**z**|**X**) is used to approximate *P* (**z**|**X**). To minimize the distance between *Q*(**z**|**X**) and *P* (**z**|**X**), the Kullback–Leibler (KL) divergence between them is minimized in a standard VAE [32] as

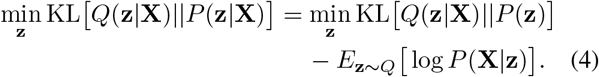

Moving to the inverse path in Fig. 1, the goal was to map **z** from **y**. The AE-Decoder (a) mapper estimated *P* (**z**|**y**), whereas the VAE-Decoder (c) and the VAE-Sampler-Decoder (d) mappers estimated *Q*(**z**|**y**). Subsequently, these three models utilized the saved *P* (**X**|**z**) decoder, from the forward training, to complete their reverse training. The Direct Convolutional model (b) trained from **y** *→* **z** *→* **X** directly in a single step, without using a saved pre-trained *P* (**X**|**z**) decoder. Finally, since the Direct Variational model (e) also trained in a single step, the optimization objective for the model became

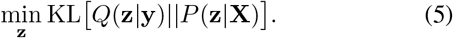

In accordance with the right-hand side of (4), (5) may be rewritten to approximate

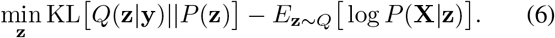

### E. Model Training and Testing

Corresponding to the models discussed in Section II-D, the family of deep networks developed and compared for this inverse imaging task were instantiated in terms of forward and inverse training paths, as seen in Fig. 2. The forward training paths for each architecture are indicated in the figure with red arrows, and bold letters beneath the arrows identify architectures that follow that path, while the reverse training paths are shown with blue arrows and italicized letters beneath the arrows. A bold italicized letter indicates that both the forward and reverse training paths for the specific model take the same route. Each box represents a particular component of the neural network. The numbers above the arrows represent the dimensions of the variables moving between two blocks.

**Fig. 2.**
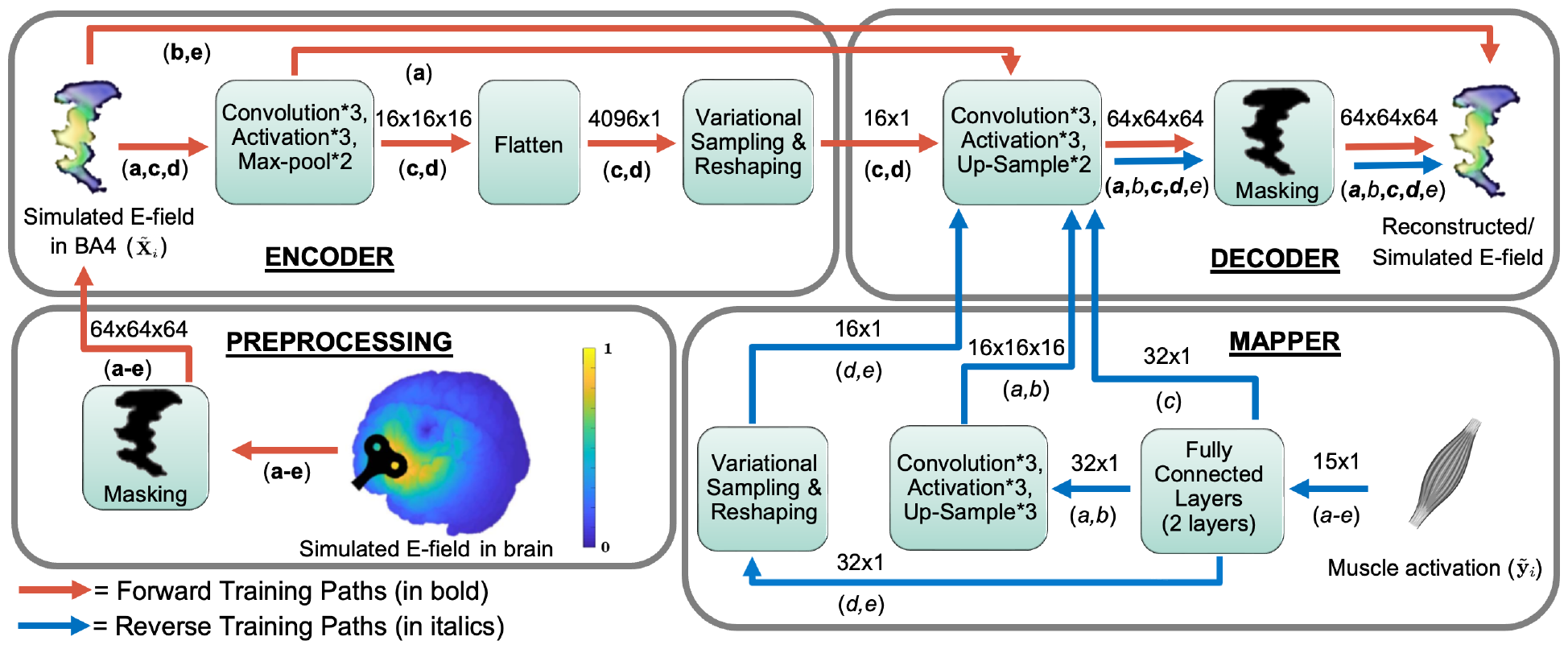
System block diagram for training. The different models were first trained along the forward paths (red arrows), and then along the reverse paths (blue arrows), as applicable. The forward training starts in the preprocessing block, continues to the encoder block and then ends in the decoder block. The reverse training begins in the mapper block, and then finishes in the decoder block. The inference paths are the same as the reverse training paths. The numbers above the arrows indicate the dimensions of the variables moving between any two blocks, while the letters (bold or italicized) below refer to the structures of each of the five architectures according to the following explanation. A bold, italicized letter indicates that both the forward and reverse training paths for a model take the same route. A number shown following an asterisk (e.g. ^*^3) indicates the number of times a layer is present inside a particular block. The five architectures are denoted as: (a) AE-Decoder, (b) Direct Convolutional, (c) VAE-Decoder, (d) VAE-Sampler-Decoder, (e) Direct Variational.

The forward training paths (**a-e**) begin with the preprocessing block (lower left) in Fig. 2. Coil parameters are chosen at random from a training set of TMS stimulations and the corresponding E-field distribution in the chosen BA4 area is estimated using the finite element simulation. A subject-specific Brodmann area 4 (BA4) binary motor mask is then applied to this E-field distribution. The resulting simulated E-field inside this mask is used as the input 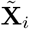 to the rest of the training network. For the AE-Decoder model (**a**), the forward path consists of three convolution and activation layers in both the encoder and the decoder blocks, with two max-pool layers in between the convolutional layers in the encoder and two upsample layers correspondingly positioned in the decoder. For the Direct Convolutional (**b**) and Direct Variational models (**e**), the forward paths directly copy over the input simulations to the reconstruction, skipping the encoder and decoder blocks entirely. For the VAE-Decoder (**c**) and VAE-Sampler-Decoder models (**d**), the encoder forward paths consist of an additional flattening and variational sampling and reshaping layer. After training along the forward paths is complete, the weights of the decoders are fixed and are not updated further.

After the forward training concludes, we begin the reverse training. The reverse training paths (*a-e*) begin with a muscle activation vector **y**_*i*_, in the mapper block (lower right) of Fig. 2. Model *(c)* passes through just the fully connected layers in the mapper block, joins the trained decoder, and completes the rest of the path as indicated in the figure. Models *(a,b)* pass through additional sets of convolution, activation, and max-pool layers in the mapper block, before being fed to the trained decoder. Finally, models *(d,e)* travel through an additional variational sampling layer, before completing similar paths through the decoder as in the forward training. Once the training on the reverse paths was complete, the weights in the mapper blocks were also fixed and the networks were ready for inference.

During training, the inputs were processed in mini-batches of size 8. Adadelta [33] was chosen as the optimizer for all models, with a learning rate of 1 to start. Model weights were saved every epoch until the training loss plateaued. If training loss plateaued for five epochs, a dynamic learning rate scheduler multiplied the learning rate by a factor of 0.7. After 20 epochs of no improvement, by at least a factor of 10^*−*5^ in the relevant loss function, the training was stopped. Training, evaluation, and output visualization of all models were done on a workstation equipped with a 9th generation Intel Core-i7 3.6 GHz CPU, 64 GB of RAM, and an NVIDIA RTX 2080 Ti GPU hardware. The programming platform used was Python 3.7.4 within JupyterLab, with support from major libraries such as TensorFlow 2.0.0, scikit-learn 0.23.2, matplotlib 3.3.2, etc. The software used for the visualization of the results was MATLAB R2019b. The source code for this work is open-source and available to the public. ^1^

During each *j*-th stimulation in model testing, a muscle activation vector 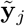 is chosen at random from a set of muscle activation vector test samples (a set separate from the samples used in training, details of which are outlined in Section II-F) is fed as an input to each inference path (the same as each reverse training path), as seen in Fig. 2. The various models follow the paths indicated by the blue arrows in the figure, using the fully trained mapper and decoder. The output from the inference path then produced an estimate of the three-dimensional E-field reconstruction 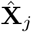, corresponding to the MEP test sample 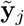.

### F. Model Parameters

The entire set of input-output data for each subject was divided into train and test sets in a 10-fold outer cross-validation (CV) arrangement. This division was stratified such that the distribution of the stimulations from the different levels of stimulation intensity (%RMT), as present in the original data set for a given subject, was preserved between individual train and test sets. The number of channels for the convolutional filters and the value of the 𝓁_1_ regularization parameter was determined by following [12]. For choosing new parameters, such as the length of the variational sampling layer or the final activation function, we took the training data portion (of subject 1 only) of one of the original folds and subdivided into a second 10-fold “train-validate” CV, where 9*/*10’ths of each fold was used to train and the last 1*/*10’th was used as validation. The lowest normalized root mean square error (NRMSE) performance across these 10 validation sets determined the best choice for tuning the relevant parameters. Once tuning parameters were fixed, all models were trained on the entire applicable training set, in each CV fold.

Referring to Fig. 2, there are two sets of convolution-activation-maxpool layers followed by a single convolutional layer in the encoder and two sets of convolution-activation-upsample layers followed by a single convolutional layer in the decoder. The number of channels in the first two convolutional layers of the encoder was 32 and 64, respectively, while for the decoder it was 64 and 32, respectively. The first two convolutional layers in both the encoder and the decoder had 3*×*3*×*3 filters, while the last one had a 1*×*1*×*1 filter and a single channel, The padding used was 1 element on each side, and the stride was 1.

Activation functions followed all convolutional and fully connected layers. The rectified linear unit (ReLU) was chosen as the activation function for all intermediate layers. The final activation function in the decoder was a bounded ReLU, which implemented the minimum value between 1 and a ReLU output [34]. This choice arose from the need to constrain the output between 0 and 1, to match the min-max scaling applied earlier at the input, and maintain the physiological interpretability of the reconstructed E-fields. Although a sigmoid activation serves the same purpose, the bounded ReLU consistently outperformed the sigmoid in the inner CV experiments we conducted and was thus used in all models in this work.

Max-pooling and upsampling layers were used to reduce and increase the sizes of the representations, respectively, and had filter windows of size 2*×*2*×*2 and a stride of 1.

All voxels not part of the BA4 motor cortex volume (∼98% of the 64×64×64 box in Fig. 2, for all three subjects) were set to zero. This was represented as ‘masking’. Although the E-field distribution itself was smooth, the resulting volume then became sparse. This allowed us to use an *𝓁*_1_ penalty, since it has been shown to be effective in convolutional sparse coding [35]. We retained an *𝓁*_1_ regularization value of 10^*−*4^, what we empirically determined earlier in [12].

Batch normalization layers were used following the convolutional layers in the mapper to prevent internal covariance shifts in the data. In our inner CV experiments, we noted that the batch normalization interestingly only improved accuracy in the first two, purely convolutional architectures, and not the variational ones, and was thus used only in models (a-b) to reduce the complexity of the variational models (c-e).

### G. Loss Functions

Each of the five networks was trained to optimize a relevant cost function. To train both the forward and reverse paths for the AE-decoder (a) and Direct Convolutional models (b), as well as the reverse path only for the VAE-decoder (c), we minimized the mean squared error (MSE) loss, which we denote as *L*_1_(*θ*), between the ground truth (GT) E-field distribution and its reconstruction, for *N* training samples as

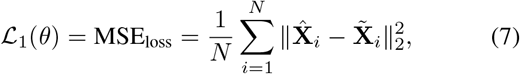

where ∥.∥_2_ denotes the Euclidean norm.

To train the forward path only for the VAE-Decoder (c) and both the forward and reverse paths for the VAE-Sampler-Decoder (d), the relevant objective was to minimize a combination of the MSE loss and the KL divergence loss from (4), given by

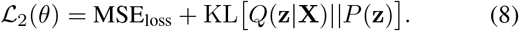

We assume the latent distribution and the surrogate to be Gaussians, parameterized as *P* (**z**) = *𝒩* (0, 1) and *Q*(**z**|**X**) = *𝒩* (*µ*(**X**), Σ(**X**)) [32]. This loss can then be rewritten as

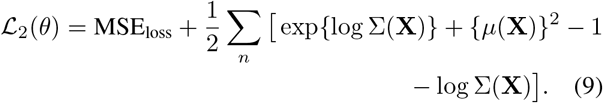

Finally, to train both the forward and reverse paths for the Direct Variational model (e), an objective consisting of the MSE loss and the relevant expression for the KL divergence loss from (6) was minimized:

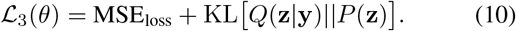

Assuming *Q*(**z**|**y**) to be a Gaussian parameterized by *𝒩* (*µ*(**y**), Σ(**y**)), (10) can be reformulated as

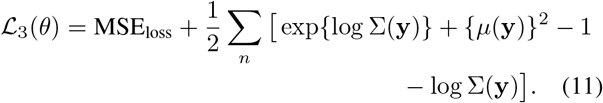

### H. Evaluation Metrics

All five models were first trained on each fold’s training set and then evaluated on the corresponding test set for each CV fold. In each CV round, the model weights were first cleared and then randomly initialized for a new iteration of training. The performance of each model was assessed on each stimulation for each of the three subjects, using three evaluation criteria. NRMSE, the primary metric of performance assessment, was calculated f the *j*-th test stimulation as

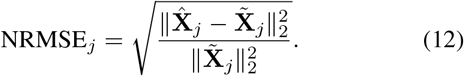

To measure the similarity between the individual reconstructions of the E-field and the respective GT, *R*^2^ was calculated as a secondary metric:

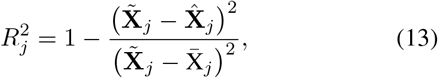

where 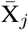 is the mean of all voxel intensities (*v*_*jk*_) contained in 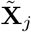 for voxels *k*, within the volume of the motor cortex **K** Finally, the center of gravity (CoG), a common outcome used in TMS mapping [2], for both the GTs and the predictions of the E-field distributions wer alculated as

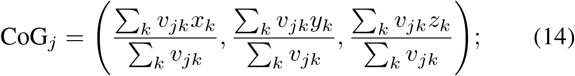

where *x*_*k*_, *y*_*k*_, and *z*_*k*_ are the Cartesian coordinates *∀k ∈* **K**. The error in CoG (CoG_error_) in the reconstructions then formed the tertiary metric:

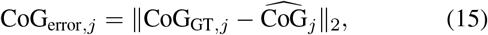

where CoG_GT,*j*_ is the GT COG for **X**_*j*_ and 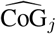 is the CoG calculated for 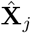 .

## III. Results

All subjects tolerated stimulation well, and no adverse events were reported. FDI resting motor thresholds for the three subjects were 50, 42, 41% maximum stimulator output respectively.

### A. Performance Across Models

Table II reports our statistics from the performance comparison of the presented models, across ten cross-validation folds, for all three subjects. The mean NRMSE and *R*^2^ are reported, across all stimulations for each subject, along with the corresponding standard errors of the mean (SEMs) calculated for a 95% level of confidence. For the first two purely convolutional models, we observed that the Direct Convolutional (b) consistently outperformed the AE-Decoder (a) across all subjects. For the next three models (c,d,e) involving VI, it was noticeable that all of them performed better than their purely convolutional counterparts. Finally, the Direct Variational model (e) consistently performed the best, both in terms of NRMSE and *R*^2^, with the VAE-Sampler-Decoder (d) as a close second.

**TABLE II.** Performance of the different models, for each subject. The means and standard errors of the means across the cross-validation folds are reported for the NRMSE and *R*^*2*^ .

| Subject | Model | NRMSE | $R^2$ |
| --- | --- | --- | --- |
| 1 | AE-Decoder | 0.294±0.014 | 0.836±0.011 |
|  | Direct Convolutional | 0.219±0.011 | 0.832±0.010 |
|  | VAE-Decoder | 0.208±0.010 | 0.852±0.011 |
|  | VAE-Sampler-Decoder | 0.184±0.008 | 0.854±0.010 |
|  | Direct Variational | **0.181±0.008** | **0.855±0.010** |
| 2 | AE-Decoder | 0.402±0.011 | 0.692±0.009 |
|  | Direct Convolutional | 0.252±0.007 | 0.720±0.009 |
|  | VAE-Decoder | 0.245±0.006 | 0.744±0.011 |
|  | VAE-Sampler-Decoder | 0.220±0.005 | 0.754±0.010 |
|  | Direct Variational | **0.213±0.006** | **0.756±0.010** |
| 3 | AE-Decoder | 0.250±0.008 | 0.566±0.012 |
|  | Direct Convolutional | 0.222±0.007 | 0.571±0.012 |
|  | VAE-Decoder | 0.216±0.007 | 0.595±0.014 |
|  | VAE-Sampler-Decoder | 0.166±0.005 | 0.682±0.014 |
|  | Direct Variational | **0.156±0.005** | **0.714±0.014** |

\* The best value for each metric in each subject is shown in bold.

To obtain qualitative insight into the E-field reconstruction fidelity, the performance of the various models is illustrated in Fig. 3 for a single stimulation for subject 3 that elicited large responses in most of the muscles. We chose an example where the models would indicate strong E-field activations, as we intend to illustrate differences across the models we developed, which can be best seen in such cases. The image on the top left of Fig. 3 shows the normalized GT E-field of the chosen stimulation, with muscle activation vector (normalized as described above) in the inset bar graph. The other five panels show the five different reconstructions. In the reconstructions using the AE-Decoder (a) and the Direct Convolutional (b) architectures (first row), we observe underestimation of the intensity of the E-fields around the CoG, producing flatter intensity profiles than are present in the GT. With the VAE-Decoder (c), where the latent space was learned from the E-fields and subsequently fixed, a similar result was observed.

**Fig. 3.**
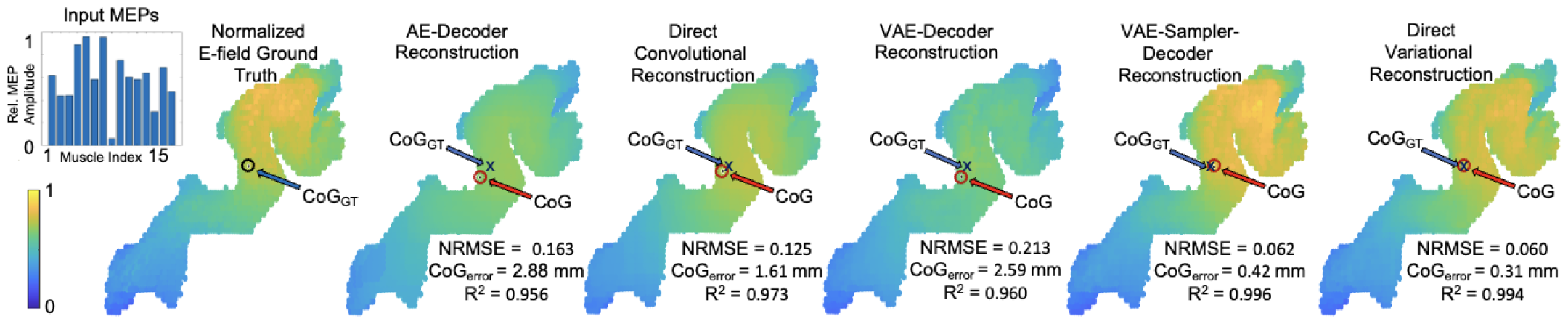
Reconstructions from the five different network architectures for subject 3, for a sample stimulation that elicited large responses in most of the muscles (the MEPs are shown as an inset on the upper left). This sample was selected in order to clearly show the differences between the various models, and thus the error values here are smaller than the averages reported in Table II. The CoG of the estimated E-field is indicated on each plot with a red circle. The range of intensities for all the maps are min-max normalized to unity, and indicated accordingly by the color bar. The NRMSE, *R*^*2*^, and CoG_error_ for each reconstruction are shown at the lower right of each map.

For the VAE-Sampler-Decoder (d) and Direct Variational (e), we observe that the reconstructions reproduced the GT E-fields with a high degree of fidelity. Although it may seem difficult to distinguish between the two outputs in this specific example, the proposed Direct Variational (e) model consistently outperformed the VAE-Sampler-Decoder (d) model in aggregate, as seen in Table II. Since the Direct Variational model provided the most accurate reconstructions, we present example results using that architecture only in the next subsections.

### B. Performance of the Direct Variational Model: Effects of Stimulation Intensity

The induced E-field is directly related to the intensity of stimulation. We therefore, analyzed the reconstruction performance of the Direct Variational model with respect to the four stimulation intensities applied. In Table III, we report the mean and SEMs (for a 95% level of confidence) of NRMSE and *R*^2^, averaged across all 10 folds for each intensity and for each subject. We did not observe a clear trend in model performance across the stimulation intensities and participants, indicating that Direct Variational model performance, in aggregate, was not sensitive to the stimulation intensity used. We recall that for subject 1 the low E-field stimulations that were discarded in preprocessing, as described in Section II-C, constituted 49 out of the 149 total for intensity of 120% RMT. That may explain why the reconstruction performance for this stimulation intensity did not match that for the other three intensities, in subject 1.

**TABLE III.** Performance of the Direct Variational model for each stimulation intensity, for each subject. The means and standard errors of the means across the cross-validation folds are reported for the NRMSE and *R*^*2*^ .

| Subject | % of RMT | NRMSE | $R^2$ |
| --- | --- | --- | --- |
| 1 | 110 | 0.172±0.015 | 0.879±0.020 |
|  | 120 | 0.213±0.033 | 0.806±0.052 |
|  | 130 | 0.176±0.019 | 0.863±0.027 |
|  | 140 | 0.176±0.021 | 0.862±0.029 |
| 2 | 110 | 0.222±0.015 | 0.770±0.039 |
|  | 120 | 0.204±0.011 | 0.792±0.035 |
|  | 130 | 0.219±0.011 | 0.743±0.043 |
|  | 140 | 0.208±0.011 | 0.717±0.042 |
| 3 | 110 | 0.169±0.014 | 0.668±0.034 |
|  | 120 | 0.152±0.012 | 0.707±0.032 |
|  | 130 | 0.136±0.011 | 0.756±0.029 |
|  | 140 | 0.166±0.013 | 0.723±0.031 |

### C. Performance of the Direct Variational Model: Effects of Muscle Response

To give insight into differences in performance across stimulations with respect to the muscle response profile, we visualize the best, average, and worst E-field reconstructions, based on NMRSE, for the Direct Variational model in Fig. 4 for the same subject as in the previous figure (subject 3). The best reconstruction, in terms of lowest NRMSE, also yielded a very low CoG_error_ and shift of the CoG (both close to zero) and a very high *R*^2^ (close to one). The reconstruction error map for this case confirms that it reproduced the ground truth E-field with very high accuracy, with a maximum normalized voxel intensity of 0.05 in the error map. We note that the mean activation of the input muscles was high across many muscles. In the stimulation with an NRMSE that was closest to the average performance for subject 3 (middle column), as reported in Table II, the NRMSE was higher, the CoG_error_ was larger, and the *R*^2^ was smaller than for the best case. This reconstruction also replicated ground truth well, though small artifacts are visible along the edges of the error map (bottom row), with a maximum normalized voxel intensity error of 0.18. The number of activated muscles and the mean activation across muscles was lower in comparison to that for the best case reconstruction. Finally, the worst reconstruction for this subject, shown in the right-hand column, corresponded to a case where the MEP activation was localized to a single muscle with a small amplitude. The NMRSE was substantially higher than for the other two examples, the CoG_error_ was correspondingly large and *R*^2^ was low. The error map showed broad regions of high normalized voxel intensities (0.2*∼*0.4) where the errors were high, with 0.43 as the maximum normalized voxel intensity.

**Fig. 4.**
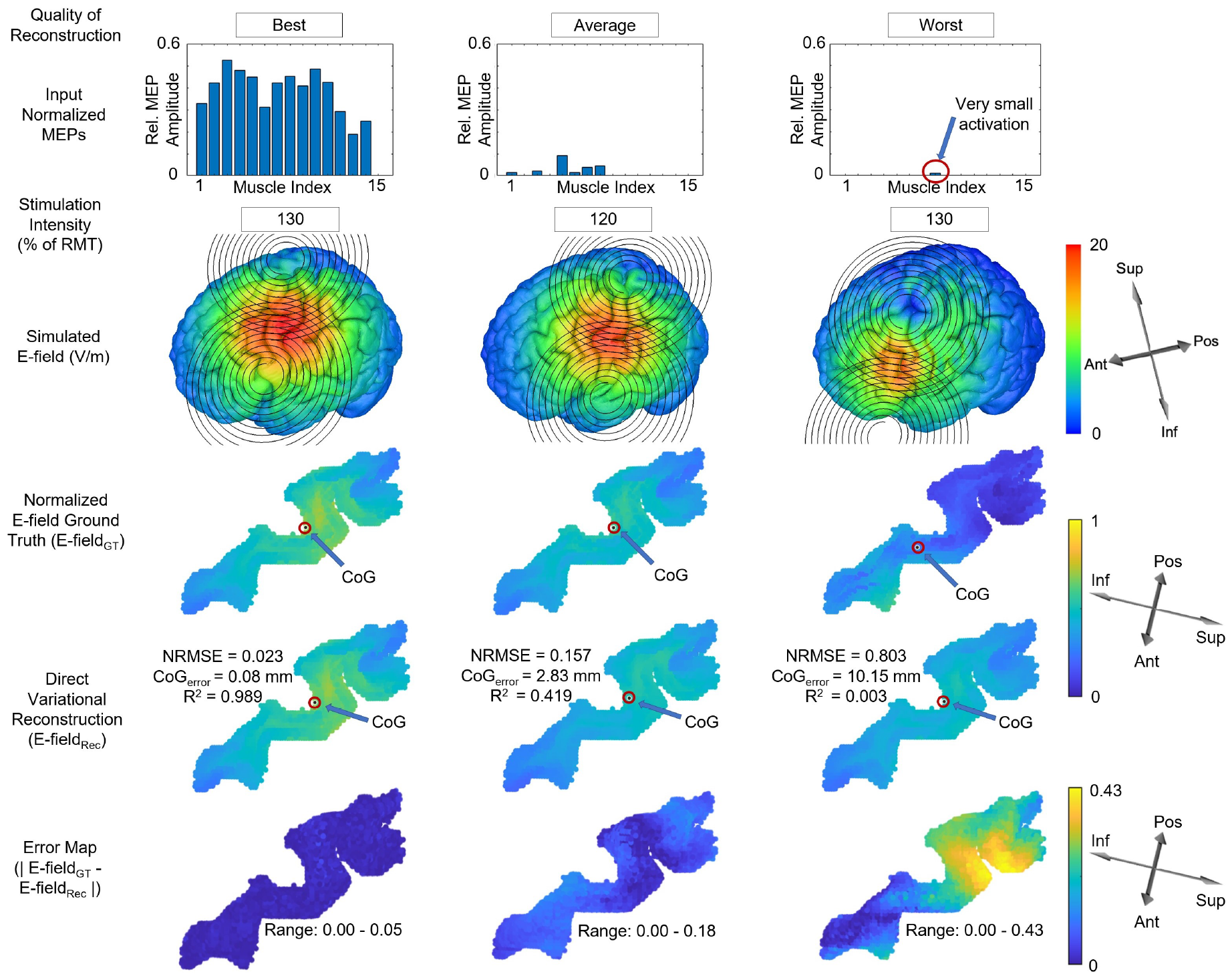
Comparison of the best, average, and worst reconstructions (in terms of NRMSE) along the columns from left to right, for subject 3, using the Direct Variational model. Starting from the top row, each column shows the normalized input MEPs, the corresponding stimulation intensity, the ground truth (GT) E-field distribution simulated on the brain, the GT E-field (E-field_GT_) on the BA4 map, the reconstructed E-field (E-field_Rec_) on the BA4, and finally an error map (|E-field_GT_ ***−*** E-field_Rec_|) on BA4 to further illustrate the accuracy of the reconstruction. The CoG is indicated on the GT and reconstructed BA4 maps with a red circle. Note that the range of intensities for all the maps are min-max scaled: the simulated E-field is shown in units of V/m, while reconstruction and error maps are normalized to unity.

To illustrate the effect of the muscle response profile on the performance of the Direct Variational model across test stimulations from all CV folds, we show scatter plots of the NRMSE against the mean (Fig. 5a) and variance (Fig. 5b) of the normalized MEPs for the same subject (subject 3). The highest error samples were largely concentrated where both the mean and variance of activation were the lowest, and NRMSE decreased with increased mean and variance of activation across muscles. As expected, the mean and variance of the muscle response to stimulation increased with increasing intensity, however, the relationship between NRMSE and intensity was variable in agreement with the aggregate data shown in Table III.

**Fig. 5.**
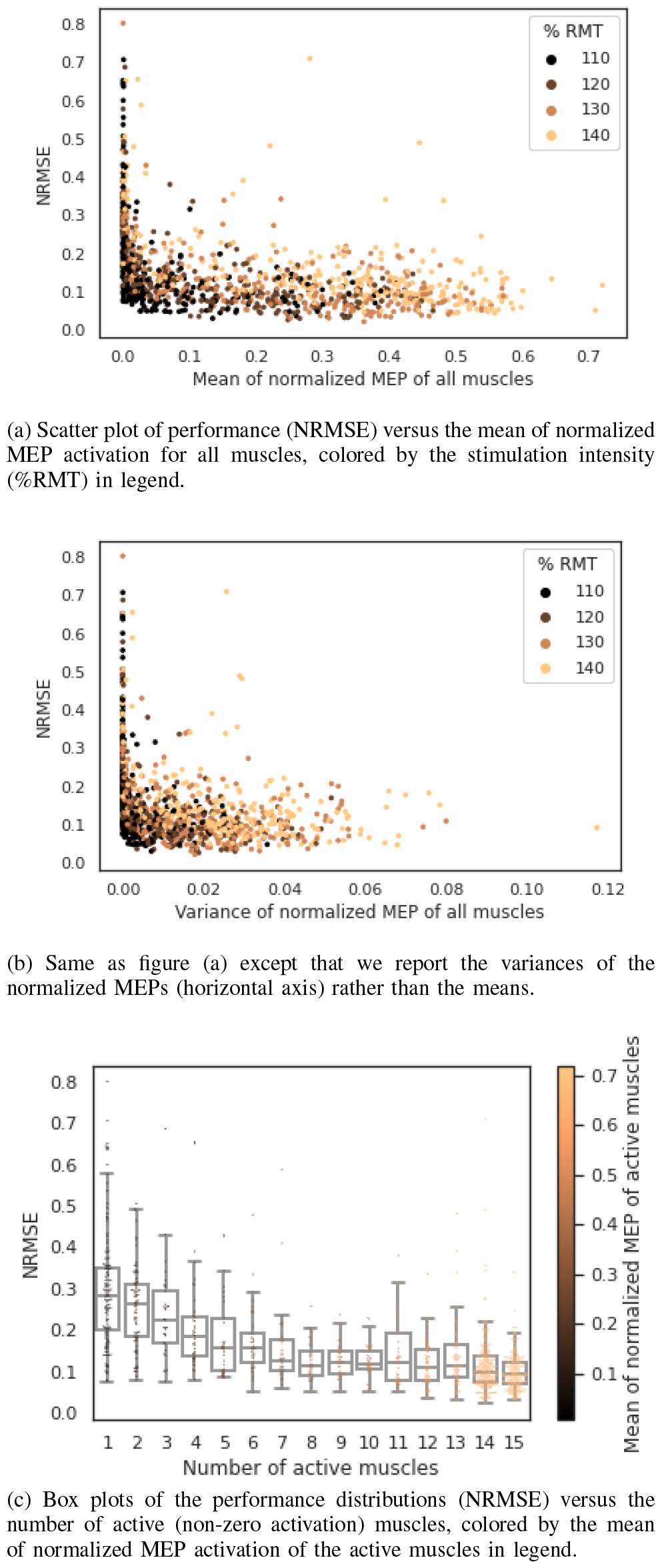
Reconstruction performance (NRMSE) for all 10 test CV fold stimulations and sensitivity to MEP mean and variance, for the Direct Variational model for subject 3.

To view the effect of the muscle response profile from a different perspective, we show the NRMSE distributions as box-plots (Fig. 5c) against the number of active (non-zero activation) muscles for test stimulations from all CV folds. As seen from the box plots, the median NRMSE tended to decrease with an increasing number of active muscles, up to about eight active muscles, beyond which it somewhat plateaued. NRMSE was notably higher for stimulations that either activated a single muscle (Fig. 5c), or produced a response *<*0.02 in normalized MEP mean of all muscles (Fig. 5a), indicating reconstructions using the Direct Variational model must be interpreted with caution for these types of stimulations.

## IV. Discussion

### A. Variational Modeling

Table II provided evidence that the purely convolutional AE structure [11], [12] and the two-stage training strategy [16], might not bring additional benefits within our current experimental framework. The VAE-Sampler-Decoder and Direct Variational models outperformed the VAE-Decoder. We speculate that this may be because VAE-Decoder did not utilize the benefits of VI in the reverse training path. The subtle difference between the outputs VAE-Sampler-Decoder and Direct Variational models could be due to the fact that the VAE-Sampler-Decoder attempted to match the **z** obtained from the *Q*(**z**|**y**) mapper with the underlying **z** (obtained from the *Q*(**z**|**X**) encoder) forming the saved *P* (**X**|**z**) decoder, whereas the proposed Direct Variational method directly optimized *P* (**X**|**z**) from samples obtained from *Q*(**z**|**y**). Our finding is thus consistent with [17], where the CNN model learned to map the sensor domain data to the image domain information using a single-stage training strategy.

A potential question may arise with regard to the choice of MSE in the loss functions of all the models, compared to the choice of NRMSE as an evaluation metric. For a regression model with a Gaussian distributed noise, as outlined in our earlier works on the forward modeling [11], [12], it can be shown that following a Bayesian approach, maximizing the log-likelihood function for the target variable is equivalent to minimizing the mean squared error (MSE) [36], [37]. So MSE, with a simple first-order derivative, was a natural choice for the loss function. While NRMSE has no such straightforward mathematical formulation to be used as a neural net loss function, it is a popular choice as an evaluation metric since it can overcome scale-dependency [36]. So one might interpret our result as saying that despite the bias in the loss function from using MSE, we still achieve reasonable performance as measured by NMRSE.

### B. Sparse and Zero Activations

In Figs. 5 & 6, we observed that the E-field reconstruction accuracy is affected by the amplitude profile of the MEPs used as the input to the inverse mapper. The E-field reconstruction was notably better for MEP vectors with larger mean amplitude, variance, and number of muscles with non-zero amplitude. Interestingly, reconstruction error was worse overall but also more variable for stimulations in which only one or two muscles were activated (had non-zero MEP amplitude). As is shown in the “worst” reconstruction (right column) example in Fig 5, single (or few) muscle stimulation can result when the coil is distant from the canonical hand area (’hand knob’) of the motor cortex or when the E-field was relatively low in amplitude and distributed. It is therefore unsurprising that it was challenging for the network to estimate the specific E-field distribution, and it ended up returning a low-intensity distributed profile for these types of stimulations. Such low amplitude MEP response profiles are proximal to the zero activation MEP profile in the vector space. Thus, the set of possible E-field distributions that can produce such MEP responses are in the vicinity of the null space of the transformation matrix equivalent of the cortico-motor mapping for the hand knob area.

In principle, it might appear beneficial to exclude a certain number of voxels in the cortex from the localization where the electric field was high but did not yield any behavioral response. For this relevant exploration, we re-ran the Direct Variational model for subject-3 with zero MEP stimulations. We observed that most of the highly activated voxels in the ground truths of such zero MEP inputs were also present in the ground truth of large response stimulation of Fig. 4. If we were to discard these highly active zero MEP ROIs, we would end up excluding patches of voxels in different regions of the reconstruction maps. Consequently, the reconstruction maps would appear unnatural and physiologically less meaningful, and also affect our performance metrics (e.g. NRMSE, R^2^) adversely during model evaluation. Thus, we did not see any particular advantage in practice for choosing to exclude such ROIs, while doing so would require substantial modification to the current study design and evaluation metrics. Future work may build upon training appropriately M2M-InvNet with zero and small MEP inputs, to improve E-field estimates for stimuli that induce sparse muscle activation.

### C. Neuroscience Interpretation

The advancement of biophysical modeling of the induced E-field generated by TMS has precipitated efforts to move past ascribing muscle activations induced by TMS to a single point on the scalp. Recently, there have been several concerted efforts to link resulting muscle activation to the complex spatial distribution of the induced electric field [9], [10], [38]. These techniques generally link the mapping between E-fields and single muscle activations, by utilizing a single function such as the log-sigmoid non-linearity. While this approach can be applied to multiple muscles, it is not suitable for the inverse mapping of multi-muscle activations. Our approach, by contrast, is capable of mapping the activation of multiple muscles simultaneously. Moreover, we utilize cascaded nonlinear activations (rectified linear units) of a neural network for our mapping, which is better suited for efficiently approximating nonlinear functions [39], even when they are smooth [40]. The generative quality of our approach, the ability to generate a high-dimensional E-field distribution for a novel muscle activation vector, may enable new investigations into the organization of muscle modules on the cortex. The mapping of multi-muscle modules on the cortex is more in line with modern representations of the motor humunculous [41] and the general idea of a mosaic representation of muscle topographies [42]. The proposed model could also prove useful in efforts to optimize coil position and mapping efficiency [43] by generating a probabilistically likely E-field distribution for activating a muscle or muscles of interest. However, we need to exercise caution in interpreting the results as the E-field distribution predicted by the model might not be actually producible with a conventional TMS coil, since the results originate from a model fit and thus may not be physically realizeable by a given coil.

### D. Limitation

There are several limitations of this work. For one, the E-fields we attempted to reconstruct from the MEPs were themselves simulated and calculated using numerical procedures from the coil position and orientation parameters. Thus it would be useful to add to the current procedure an additional step in the inverse calculation that tries to reconstruct the coil parameters. This is a subject of our future work. Reconstruction results may have potentially been influenced by EMG cross-talk in the recorded MEPs. Common cross-correlation techniques for assessing cross-talk in voluntary EMG are not suited for the assessment of cross-talk in evoked potentials. A small number of investigations have utilized different approaches to assess cross-talk in MEPs with widely varying results. The ability to discern physiological co-activation (via possible synergy mechanisms) from EMG cross-talk is an important area of research beyond the scope of the work presented here, but would likely benefit the ability to accurately reconstruct cortical topographies associated with multi-muscle activations. Stimulus intensity, pulse shape (monophasic/biphasic) and current direction (PA/AP) are known to influence MEPs [44]. In this study, a monophasic stimulation waveform and PA current direction were selected based on common parameters used in TMS mapping and the availability of equipment for stimulation. It is unknown whether these parameters influenced the reconstruction quality of our model. Additional research is needed to assess the effect of stimulus intensity, pulse shape, and current direction on inverse mapping of motor topography using TMS. The data set constituted only three subjects. Results from more subjects are needed to validate the robustness of the proposed model. MEPs from clinical patients often tend to be smaller and more sparse compared to healthy controls. Tuning of the M2M-InvNet structure and parameters may be required for greater utility in this population. Finally, although we tested five different CNN architectures, there may be yet another architecture that would perform even better.

### E. Future Work

In future work, we plan to include cortical motor topography mapping using active learning [24] and to study the generation of the volume conductor model by deep learning, to determine if we can combine these with the current expert user-guided mapping and the segmentation-finite element simulation pipeline, respectively, or perhaps even replace either or both entirely. As we obtain more experience with our current approach, we may be able to develop generic or semi-personalized models without the need for subject-specific volume conduction models, which could broaden applicability. Aside, studying in detail the optimal number *m*_*o*_ of muscles to measure and, which muscles to choose, is another interesting but involved problem that could be an interesting topic for future work.

## V. Conclusion

In this work, five 3D CNN models were systematically designed to estimate TMS-induced E-field distributions on the BA4 motor cortex from resultant muscle activation measured as MEPs in an inverse imaging task. Our Direct Variational generative model, which directly optimized the latent space from both the MEP input and the E-field output during training, emerged as the best performing model, and thus our candidate of choice for M2M-InvNet. In particular the Direct Variational performed better than our other four models on all three metrics of evaluation; it showed the lowest root mean square error, the highest average fidelity reconstruction, and the smallest average shift in the center of gravity of the induced fields, when compared to the ground truth. Subsequent examination of M2M-InvNet inference at different levels of stimulation intensity revealed that both the location and intensity of the stimulation in the target area had substantial impacts on the reconstruction performance, and the number of muscles activated and the mean and variance of their MEPs all generally correlated positively (up to a threshold, with the number of active muscles) with performance.

## Footnotes

1 https://github.com/neu-spiral/TMS-EMG.

